# Application of AI-Designed OpenCRISPR-1 for Highly Efficient Gene Editing in Soybean and *Nicotiana benthamiana*

**DOI:** 10.64898/2026.08.27.747635

**Authors:** Cuong X. Nguyen, Phat T. Do, Thu Minh Tran

**Author notes:** Corresponding author **Cuong X. Nguyen**.

## Abstract

The widespread application of CRISPR/Cas genome editing for commercial crop improvement is currently hindered by a complex and restrictive intellectual property (IP) landscape. The recent development of OpenCRISPR-1, a fully AI-designed and open-source Cas9-like nuclease, provides a promising, IP-unencumbered alternative; however, its efficacy in dicotyledonous plants remains largely uncharacterized. Here, we report the successful adaptation of the OpenCRISPR-1 system for highly efficient targeted mutagenesis in dicots. We constructed a plant-optimized binary vector (pBSE-OpenCRISPR-1) and validated its editing capability across two species. In soybean (*Glycine max*), targeting the *GmFAD2-1B* gene via an *Agrobacterium rhizogenes*-mediated hairy root transformation system yielded a robust mutation rate of approximately 50%. In *Nicotiana benthamiana*, stable *Agrobacterium*-mediated transformation targeting the phytoene desaturase homologs (*NbPDSa/b*) achieved a 75% editing efficiency in T0 lines, with up to 13.8% of events displaying complete homozygous or biallelic mutations and the corresponding visible albino phenotypes. Deep amplicon and Sanger sequencing revealed a characteristic mutation profile dominated by 1-bp insertions and small deletions occurring two to three nucleotides upstream of the PAM. These results demonstrate that the AI-designed OpenCRISPR-1 system is a highly active and versatile nuclease for dicot genome engineering, offering a powerful, commercially unencumbered tool to accelerate global crop trait improvement.

**Key message:** The AI-designed, open-source OpenCRISPR-1 system mediates highly efficient targeted mutagenesis in dicotyledonous plants, achieving up to 75% editing efficiency and robust biallelic mutations in T0 transformants, thereby providing a powerful, IP-unencumbered platform for agricultural biotechnology.

## Introduction

CRISPR/Cas systems have revolutionized plant biology and agricultural biotechnology. By enabling precise, targeted modifications within plant genomes, the CRISPR/Cas system has accelerated the functional characterization of plant genes and facilitated the rapid development of improved crop varieties (Tuncel et al., 2025). Among the diverse array of nucleases, the *Streptococcus pyogenes* Cas9 (SpCas9) has emerged as the most widely adopted tool for plant genome engineering due to its robust activity and simplicity. Despite its widespread success, the complex and restrictive intellectual property landscape surrounding SpCas9 and Cas12a poses a major hurdle for the commercialization of edited crops, especially in developing countries (Chaurasia, 2024; Jefferson et al., 2021; Kim et al., 2024; Lin et al., 2025). Consequently, there is a substantial demand for novel, highly efficient, and IP-unencumbered genome editing tools to democratize crop improvement.

Recent breakthroughs in generative artificial intelligence (AI) and deep learning have opened new avenues for protein engineering, enabling the *de novo* design of programmable nucleases. Large language models (LLMs) trained on vast datasets of natural diverse CRISPR-Cas systems can now generate synthetic nucleases that do not exist in nature. Recently, Ruffolo et al.(Ruffolo et al., 2025) reported the development of OpenCRISPR-1, a fully AI-designed Cas9-like nuclease. In mammalian cell cultures, OpenCRISPR-1 demonstrated editing efficiencies comparable to or exceeding those of SpCas9, while exhibiting a significantly reduced off-target profile and requiring a standard NGG PAM. Furthermore, the system has proven highly versatile, having been successfully adapted for high-fidelity base and prime editing in mammalian systems (Hwang et al., 2026; Ruffolo et al., 2025). Crucially, OpenCRISPR-1 was released under an open-source licensing framework, effectively eliminating commercial royalties and IP barriers (Ruffolo et al., 2025).

Following its success in mammalian systems, the application of OpenCRISPR-1 is just beginning to expand into agricultural biotechnology. Recently, a plant-optimized OpenCRISPR-1 was adapted to mediate robust gene knockout, base editing, and prime editing in the monocot crop rice (*Oryza sativa*) (Das et al., 2026; Gupta et al., 2026). The optimized OpenCRISPR-1 was also applied in the alfalfa plant (*Medicago sativa*); however, the editing efficiency was relatively low (approximately 30%)(Alam et al., 2026).

Building upon these initial findings, in this study, we report the successful adaptation and application of the AI-designed OpenCRISPR-1 system for highly efficient genome engineering in dicotyledonous plants. To evaluate its efficacy, we engineered the pBSE-OpenCRISPR-1 binary vector system optimized for plant expression. We tested the targeted mutagenesis capability of OpenCRISPR-1 in two distinct dicot species. First, we targeted the *GmFAD2-1B* gene, a key regulator of fatty acid biosynthesis in soybean using an *Agrobacterium rhizogenes*-mediated hairy root transformation system for rapid validation of the system’s effectiveness. Second, we targeted the phytoene desaturase genes (*NbPDSa/b*) in the model plant *Nicotiana benthamiana* using *Agrobacterium*-mediated stable transformation. Our results demonstrate that the AI-designed OpenCRISPR-1 is a highly active and efficient nuclease in dicot plant cells, offering a powerful, commercially unencumbered alternative for global crop trait engineering.

## Materials and methods

### Vector constructions

To generate the pBSE-OpenCRISPR-1 construct, the open reading frame (ORF) of *openCRISPR-1* was flanked by an N-terminal 3×FLAG-NLS tag and a C-terminal NLS signal. Expression of the construct was driven by the *GmUBQ* promoter and terminated by the *rbcs* E9 terminator. The individual components were isolated via PCR amplification using Q5 High-Fidelity DNA Polymerase (M2499, NEB) and the primers listed in **Table S1**. Specifically, the *GmUBQ* promoter carrying the 3×FLAG-NLS signal was amplified from pFGC-GmUBQ-Cas-GW (Cuong X. Nguyen et al., 2021), the *openCRISPR-1* ORF was amplified from Addgene plasmid #221565 (Ruffolo et al., 2025), and the *rbcs* E9 terminator was amplified from pHUE411 (Addgene plasmid #62203) (Xing et al., 2014).

The three resulting PCR fragments were assembled into a pBSE901 backbone (Addgene plasmid #91709) (Chen et al., 2017), previously digested with *Sbf*I and *Eco*RI, using the Gibson assembly approach (E2621, NEB) (Gibson et al., 2009). Following verification of the intermediate construct via whole-plasmid sequencing, a synonymous mutation was introduced using the Q5 Site-Directed Mutagenesis Kit (E0554S, NEB) to remove an internal *Bsa*I restriction site present in the *openCRISPR-1* sequence. The final pBSE-OpenCRISPR-1 construct was subsequently verified with Plasmidsaurus Whole Plasmid Sequencing (Oxford Nanopore, R10.4.1).

To construct functional OpenCRISPR-1 expression vectors for validating gene editing efficacy, DNA oligonucleotides (primer pairs) targeting the second exon of *GmFAD2-1B* (*Glyma*.*20G111000*) and the first exon of *NbPDSa/b* (*Nbe05g35010*/*Nbe06g25970*) in soybean and *N. benthamiana*, respectively (Goodstein et al., 2012) were synthesized by IDT. The oligonucleotides were annealed and ligated into the *Bsa*I-digested pBSE-OpenCRISPR-1 backbone. Positive constructs were verified by Sanger sequencing before being mobilized into *Agrobacterium* for hairy root and leaf disc transformation.

### Soybean hairy root and *Nicotiana benthamiana* stable transformation

Transgenic hairy roots of soybean (cv. Williams 82) were induced using *Agrobacterium rhizogenes*-mediated transformation, as previously described (Chen et al., 2018), with minor modifications. The binary vector pBSE-OC1-GmFAD was mobilized into *A. rhizogenes* strain K599 via electroporation and used for hairy root induction. Soybean seeds were sterilized with chlorine gas for 20–24 h, then germinated on MS medium (PlantMedia, SKU #30630067). Four to five days post-germination, wounded cotyledons were immersed in an *A. rhizogenes* suspension (OD600 = 0.6) for 30 min. The explants were transferred to co-cultivation medium and cultured for 3 days, then transferred to resting medium supplemented with 100 mg/L timentin and 250 mg/L cefotaxime for root induction and elongation. Two weeks after transformation, hairy roots were pooled and collected for genotyping.

For transformation of *N. benthamiana*, an *Agrobacterium*-mediated leaf disc approach, as previously described (Gantner et al., 2019) was utilized with minor modifications. *In vitro* leaves were excised and immersed in a suspension of *Agrobacterium tumefaciens* strain LBA4404 (OD600 = 0.5) harboring pBSE-OC1-NbPDS for 20 min. The explants were transferred to a co-cultivation medium and cultured for 2–3 days, then transferred to a selection medium supplemented with 100 mg/L timentin, 250 mg/L cefotaxime, and 3 mg/L bialaphos sodium (PlantMedia, SKU #40210002). Regenerated shoots were transferred to a rooting medium and subsequently moved to a growth chamber for further growth and analysis.

### Genomic DNA isolation, mutation analysis and amplicon sequencing

Genomic DNA from pooled soybean hairy roots and leaf tissues of *N. benthamiana* transgenic events were extracted using CTAB method (Doyle, 1990). Specific primers flanking the target sites of the candidate genes were used for PCR amplification with Q5 High-Fidelity DNA Polymerase (M2499, NEB) or DreamTaq DNA Polymerase (EP0713, Thermo Fisher Scientific). To analyze mutations via CAPS analysis, PCR products were digested with *Nco*I at 37 °C overnight and resolved on a 1.5–2% agarose gel. For genotyping by amplicon sequencing, PCR products were purified using a DNA extraction kit (K0691, Thermo Fisher Scientific) and subsequently submitted to Plasmidsaurus for Genotyping Analysis (Oxford Nanopore, R10.4.1). For Sanger Sequencing, purified PCR products were ligated into the pJET1.2 cloning system (K1232, Thermo Fisher Scientific) before being sent for sequencing. The DNA sequences derived from the putative mutants and wild-type plants were aligned using the online program Muscle (https://www.ebi.ac.uk/Tools/msa/muscle/), and MegaX (Kumar et al., 2018) to identify the mutations induced by OpenCRISPR-1.

## Results

### Development of the AI-designed OpenCRISPR-1 platform for soybean hairy root

To establish a robust platform for inducing targeted double-strand breaks (DSBs) in dicotyledonous plants, we constructed the pBSE-OpenCRISPR-1 binary vector system. This construct utilizes the AI-designed OpenCRISPR-1 nuclease (which has previously demonstrated editing activity comparable to SpCas9 at on-target sites bearing an NGG protospacer adjacent motif (PAM) (Ruffolo et al., 2025), driven by the highly active soybean ubiquitin (*GmUBQ*) promoter (Hernandez-Garcia et al., 2010; Cuong X Nguyen et al., 2021). To ensure efficient nuclear import, the *openCRISPR-1* coding sequence was flanked by nuclear localization signals (NLS) at both the N- and C-termini. These components were assembled into the pBSE901 vector backbone, which features an *AtU6-26* promoter driving the SpCas9 single guide RNA (sgRNA) scaffold and utilizes *Bsa*I restriction sites for the rapid assembly of target-specific gRNA expression cassettes into the final binary vector (Chen et al., 2017; Xing et al., 2014)**(Fig.1A)**.

To evaluate the editing activity of the pBSE-OpenCRISPR-1 system, a target sequence located in the second exon of the *GmFAD2-1B* gene, which encodes a key regulator of fatty acid biosynthesis in soybean, was designed and assessed using *Agrobacterium rhizogenes*-mediated hairy root transformation. The selected target sequence contains an endogenous *Nco*I restriction site, which was utilized to develop a cleaved amplified polymorphic sequence (CAPS) assay to rapidly assess the editing efficiency of the system (**Fig. 1B**).

**Figure 1.**
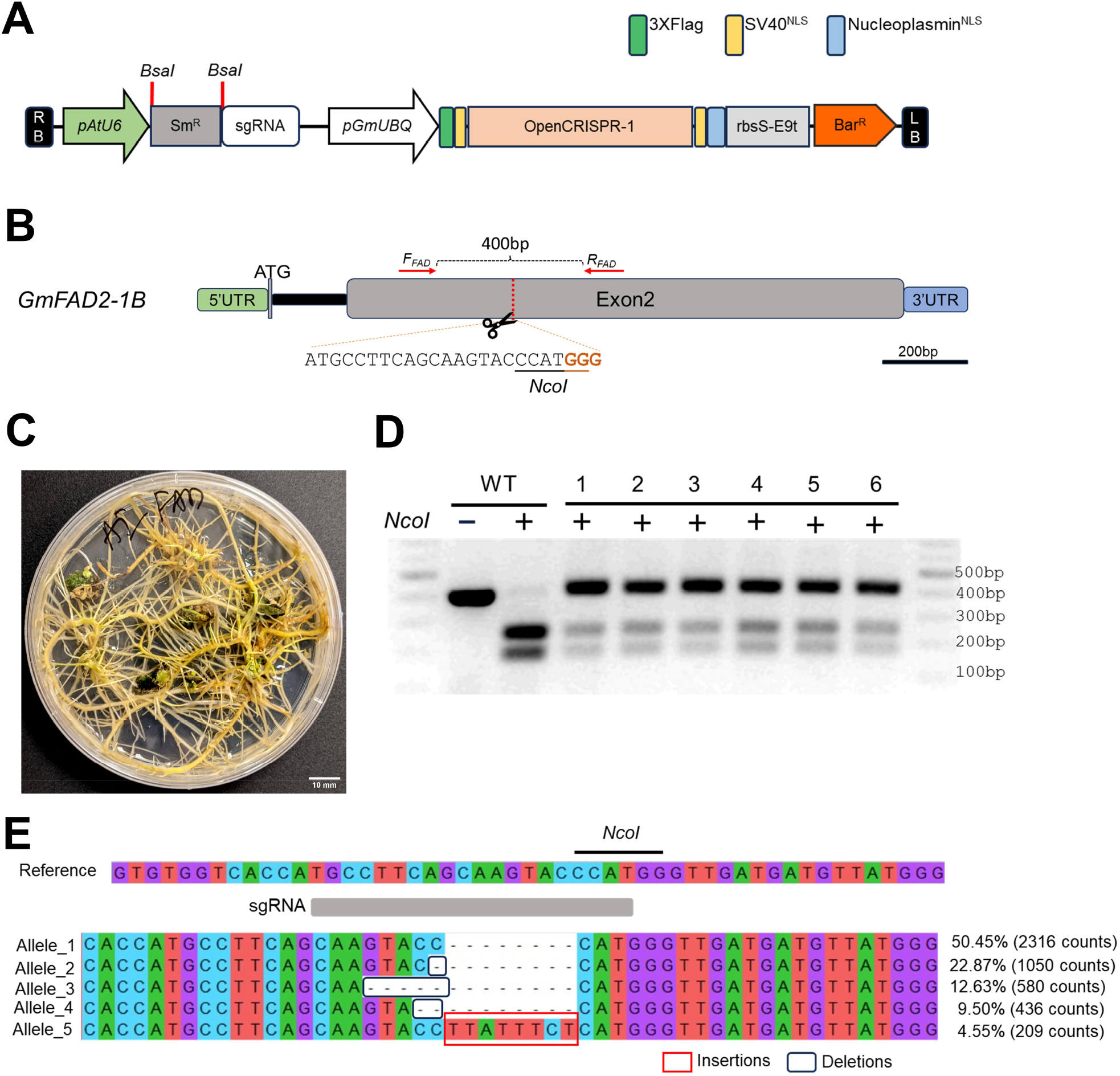
Highly efficient targeted mutagenesis in soybean using the AI-designed OpenCRISPR-1 system. **(A)** Schematic representation of the T-DNA region of the pBSE-OpenCRISPR-1 binary vector. The *openCRISPR-1* coding sequence is flanked by an N-terminal 3×FLAG tag and nuclear localization signals (NLS) at both the N- and C-termini. Nuclease expression is driven by the soybean ubiquitin promoter (*pGmUBQ*) and terminated by the *rbcs* E9 terminator (*rbsS-E9t*). The single guide RNA (sgRNA) expression cassette is driven by the *Arabidopsis U6-26* promoter (*pAtU6*), with *Bsa*I restriction sites (red lines) utilized for the rapid cloning of target-specific protospacers. **(B)** Schematic representation of the *GmFAD2-1B* gene structure and the designated OpenCRISPR-1 target site. The target sequence is located within Exon 2. The protospacer sequence is shown with the NGG protospacer adjacent motif (PAM) underlined in orange. An endogenous *Nco*I restriction site (CCATGG) overlapping the expected Cas cleavage site (dashed red line) was utilized for the cleaved amplified polymorphic sequence (CAPS) assay. Red arrows indicate the binding sites for the PCR genotyping primers. **(C)** Representative photograph of transgenic soybean hairy roots induced by *Agrobacterium rhizogenes* strain K599 harboring the pBSE-OC1-GmFAD vector, taken two weeks post-infection. Scale bar = 10 mm. **(D)** CAPS assay of the wild-type (WT) and six independent pools of transgenic soybean hairy roots. PCR amplicons spanning the target site were digested with *Nco*I. The WT amplicon is completely digested into two smaller fragments, whereas edited hairy root pools exhibit a resistant (undigested) upper band, indicating the disruption of the *Nco*I recognition site by OpenCRISPR-1-mediated mutagenesis. **(E)** Representative mutant alleles identified via deep amplicon sequencing at the *GmFAD2-1B* target locus. The wild-type reference sequence is shown at the top. Induced deletions are represented by black dashes, and an insertion is highlighted within a red box. The PAM sequence and expected cleavage site are indicated below the alignment.

Two weeks after infection, the induced hairy roots were randomly pooled and genotyped using the CAPS assay (Fig. 1C). While the target PCR fragment amplified from the wild-type sample was completely digested by *Nco*I, six independent pools of hairy roots transformed with *A. rhizogenes* harboring the pBSE-OC1-GmFAD binary vector exhibited resistant (undigested) bands following *Nco*I digestion (Fig. 1D). These results indicate that the pBSE-OpenCRISPR-1 system actively edits the endogenous *GmFAD2-1B* gene in soybean.

To further characterize the precise mutations, PCR products amplified from one of the hairy root pools were analyzed via deep amplicon sequencing using the Oxford Nanopore Technologies platform. Deep amplicon sequencing revealed a robust mutation rate of 49.55% at the target site. The mutations were predominantly characterized by a high frequency of deletions (45%), which consistently occurred three nucleotides upstream of the protospacer adjacent motif (PAM) (Fig. 1E). Additionally, Sanger sequencing of the resistant (undigested) amplicons confirmed the presence of small insertion and deletion (indel) mutations occurring two to three nucleotides upstream of the PAM (Fig. S1). Collectively, these data demonstrate that OpenCRISPR-1 provides a highly efficient, open-source platform for DSB-mediated, site-specific mutagenesis in soybean.

### High-frequency editing in stable *Nicotiana benthamiana* transformants

In addition to the rapid assessment approach using hairy root transformation in soybean, the editing performance of the OpenCRISPR-1 system was evaluated in stably transformed *N. benthamiana*. Phytoene desaturase (PDS) plays an essential role in the carotenoid biosynthesis pathway. In *N. benthamiana*, two homologs, *NbPDSa* and *NbPDSb*, encode PDS, and the simultaneous frameshift disruption of both genes results in a visible albino phenotype (Komatsu et al., 2020). A conserved sequence present in the first exon of both *NbPDS* genes, which contains an endogenous *Nco*I restriction site, was identified and utilized for targeted mutagenesis in *N. benthamiana* using the OpenCRISPR-1 system (Fig. 2A). The pBSE-OC1-NbPDS vector was delivered into *N. benthamiana* via an *Agrobacterium tumefaciens*-mediated transformation approach. Thirty-six independent transgenic events were regenerated on selection medium supplemented with bialaphos and were subsequently analyzed to assess the editing efficacy of the OpenCRISPR-1 system.

**Figure 2.**
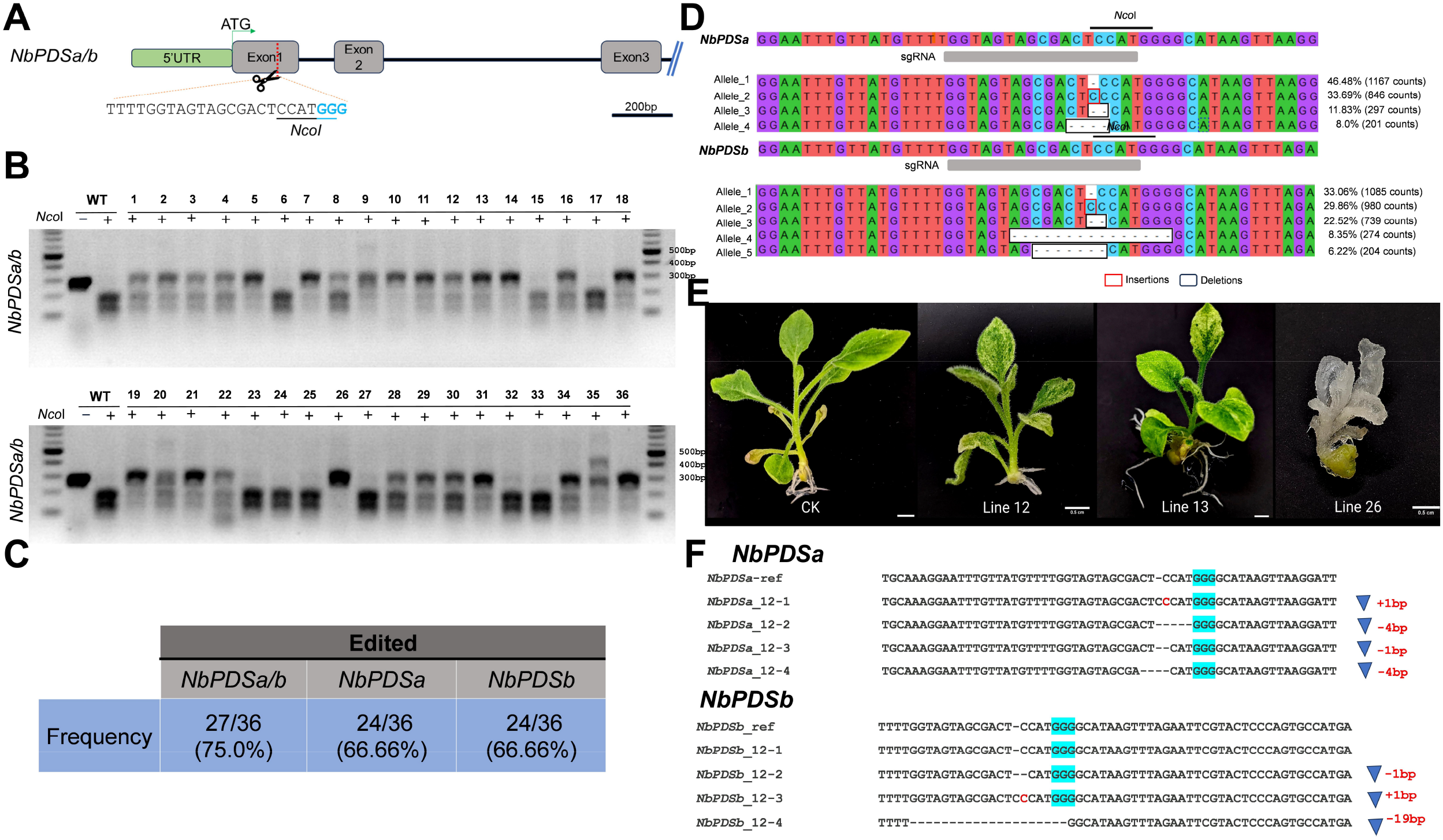
High-frequency targeted mutagenesis of *NbPDS* genes in stable *Nicotiana benthamiana* transformants using OpenCRISPR-1. **(A)** Schematic representation of the *NbPDSa/b* gene structure and the designated OpenCRISPR-1 target site. The target sequence is located within Exon 1. The NGG protospacer adjacent motif (PAM) is highlighted in blue. An endogenous *Nco*I restriction site (CCATGG) overlapping the expected Cas cleavage site (red dashed line with scissors) is underlined and was utilized for the cleaved amplified polymorphic sequence (CAPS) assay. **(B)** CAPS assay of wild-type (WT) and 36 independent T0 transgenic *N. benthamiana* events. PCR amplicons flanking the target sites of both *NbPDSa* and *NbPDSb* were digested with *Nco*I (+). The WT amplicon is completely digested, whereas edited lines exhibit higher-molecular-weight resistant (undigested) bands. Line 26 displays complete resistance to *Nco*I digestion. **(C)** Summary table detailing the editing frequencies in T0 transgenic lines at the *NbPDSa/b* loci, determined via locus-specific CAPS assays. **(D)** Representative mutant alleles identified via deep amplicon sequencing at the *NbPDSa* (top) and *NbPDSb* (bottom) target loci. The wild-type reference sequence is shown at the top of each alignment. Induced insertions are highlighted within red boxes, and deletions are indicated by white boxes, along with their respective read count frequencies. **(E)** Representative phenotypes of T0 transgenic *N. benthamiana* plants. CK indicates the wild-type control exhibiting normal green development. Lines 12 and 13 display chimeric variegated phenotypes. Line 26 displays a complete albino phenotype, consistent with the simultaneous and complete disruption of both *NbPDSa* and *NbPDSb* homologous genes. Scale bars =5mm. **(F)** Sanger sequencing alignment characterizing the mosaic editing patterns in the *NbPDSa* and *NbPDSb* loci of the variegated event 12. The PAM sequence is highlighted in cyan. Specific insertion (+, red text) and deletion (-, blue dashes) mutations are quantified on the right.

To broadly evaluate the editing activity, a primer pair flanking the target sites of both the *NbPDSa* and *NbPDSb* genes was used for initial PCR genotyping. The PCR amplification results showed that while there was no visible difference in the size of the PCR products among most of the transgenic lines compared to the wild-type, event 35 possessed a distinctly larger band (indicating an estimated insertion of ∼150 bp) in addition to the expected wild-type band (Fig. S2). Subsequent CAPS analysis using *Nco*I digestion of these PCR products revealed that out of the 36 transgenic events, 27 lines exhibited resistant (undigested) bands following *Nco*I treatment, including one line (line 26) that was completely resistant to *Nco*I digestion (Fig. 2B). These results indicate that the pBSE-OpenCRISPR-1 system is highly efficient, achieving a 75% editing rate (27/36) in *N. benthamiana* and demonstrating the capacity to simultaneously mutate both homologous genes in the T0 generation

To characterize the editing efficacy for each *NbPDS* gene individually, primer pairs that specifically amplify either the *NbPDSa* or *NbPDSb* locus were designed and utilized for mutation identification (Table S1). The locus-specific CAPS assay revealed an editing rate of approximately 66.7% for each gene, with 24 out of the 36 lines exhibiting resistant (undigested) bands following *Nco*I digestion (Fig. 2C, Fig. S3). Importantly, the frequency of complete resistance to *Nco*I digestion, indicating mutations on all alleles, was five times higher for the *NbPDSb* gene compared to the *NbPDSa* gene (5 events for *NbPDSb* versus 1 event for *NbPDSa*) (Fig. S3). These results suggest that the mutation frequency generated by the pBSE-OpenCRISPR-1 system can vary across different target loci in *N. benthamiana*. Furthermore, these data demonstrate the system’s capacity to simultaneously induce complete (homozygous or biallelic) mutations at a rate of up to 13.8% (5/36) in the T0 generation.

To further characterize the precise mutations, genomic DNA from the transgenic lines was pooled and amplified using locus-specific primer pairs for either *NbPDSa* or *NbPDSb*. The resulting PCR products were subsequently analyzed via deep amplicon sequencing using the Oxford Nanopore Technologies platform. Deep amplicon sequencing revealed robust mutation rates ranging from 53.52% to 66.94% at the target sites. The mutation profiles were predominantly characterized by a high frequency of 1-bp insertions (29.86%–33.69%), followed by 1-bp deletions and other small deletions. Consistently, these mutations occurred two to three nucleotides upstream of the protospacer adjacent motif (PAM) (Fig. 2D).

Because the simultaneous disruption of both the *NbPDSa* and *NbPDSb* genes results in a visible albino phenotype, we evaluated the transgenic lines for this distinctive trait. While control plantlets exhibited healthy green development, several transgenic events displayed either complete albino or chimeric variegated phenotypes (Fig. 2E). Notably, the complete albino phenotype observed in line 26 was consistent with its complete resistance to *Nco*I digestion at both the *NbPDSa* and *NbPDSb* loci (Fig. 2B, Fig. S3). To further investigate these mosaic editing patterns, the variegated phenotypes of events 12 and 13 were selected for detailed characterization of the *NbPDSa* and *NbPDSb* genes using Sanger sequencing. The sequencing results revealed that these mutant lines possessed several small, chimeric insertion and deletion (indel) mutations (Fig. 2F, Fig. S4).

## Discussion

The rapid evolution of CRISPR/Cas technologies has transformed plant functional genomics, yet the restrictive intellectual property (IP) landscape surrounding canonical nucleases like SpCas9 continues to impede the commercialization of edited crops, particularly for public research institutions and emerging agricultural biotechnology sectors in developing nations. In this study, we demonstrated that the fully AI-designed, open-source OpenCRISPR-1 system is highly active in dicotyledonous plants, bridging a critical gap in the current plant genome editing toolkit.

While recent studies successfully deployed OpenCRISPR-1 in the monocot *Oryza sativa* (Das et al., 2026; Gupta et al., 2026), its initial application in the dicot *Medicago sativa* yielded relatively modest editing efficiencies of around 30% (Alam et al., 2026). By engineering the plant pBSE-OpenCRISPR-1 binary vector, we achieved robust editing rates of approximately 50% in soybean hairy roots targeting the *GmFAD2-1B* locus, and up to 75% in stable *N. benthamiana* transformants targeting the *NbPDSa/b* homologs. Notably, the system was capable of generating complete (homozygous or biallelic) mutations at a frequency of 13.8% in the T0 generation of *N. benthamiana*, producing the expected visible albino phenotypes.

Our deep amplicon and Sanger sequencing analyses revealed that OpenCRISPR-1 predominantly generates small (1-bp) insertions and deletions exactly two to three nucleotides upstream of the NGG PAM. This specific mutation profile is highly consistent with the established cleavage dynamics of natural SpCas9, confirming that AI-generated nucleases can precisely mimic the biochemical fidelity of their natural counterparts while maintaining a distinct, unencumbered sequence identity.

Collectively, these results establish OpenCRISPR-1 as a highly efficient and versatile nuclease for dicot genome engineering. By validating its robust targeted mutagenesis activity in both a major leguminous crop and a foundational model plant, this work significantly expands the utility of generative AI-designed proteins in agricultural biotechnology. As an open-source platform, OpenCRISPR-1 provides global researchers and breeders with a powerful, accessible tool to accelerate trait engineering, ultimately democratizing the development and commercialization of next-generation crops.

## Data Availability

All materials reported in this paper will be shared by the corresponding authors upon reasonable request.

## Acknowledgments

We thank staffs at the PCB, IB for taking care of the plants.

## Author Contributions

CXN conceived and conceptualized the study. CXN designed, performed the experiments, and analyzed data. CXN wrote the original draft manuscript with contributions from the authors. CXN, PTD and TMT revised and proof-read the manuscript. All authors have read and approved the manuscript.

